# Assessing NGS-based computational methods for predicting transcriptional regulators with query gene sets

**DOI:** 10.1101/2024.02.01.578316

**Authors:** Zeyu Lu, Xue Xiao, Qiang Zheng, Xinlei Wang, Lin Xu

**Affiliations:** Department of Statistics and Data Science, Moody School of Graduate and Advanced Studies, Southern Methodist University, Dallas, TX, USA; Quantitative Biomedical Research Center, Peter O’Donnell Jr. School of Public Health, University of Texas Southwestern Medical Center, Dallas, TX, UƒSA; Department of Mathematics, University of Texas at Arlington, Arlington, TX 76019, USA; Center for Data Science Research and Education, College of Science, University of Texas at Arlington, Arlington, TX 76019, USA; Department of Pediatrics, Division of Hematology/Oncology, University of Texas Southwestern Medical Center, Dallas, TX, USA

**Keywords:** transcriptional regulator, benchmarking, query gene set, next-generation sequencing, prediction

## Abstract

This article provides an in-depth review of computational methods for predicting transcriptional regulators with query gene sets. Identification of transcriptional regulators is of utmost importance in many biological applications, including but not limited to elucidating biological development mechanisms, identifying key disease genes, and predicting therapeutic targets. Various computational methods based on next-generation sequencing (NGS) data have been developed in the past decade, yet no systematic evaluation of NGS-based methods has been offered. We classified these methods into two categories based on shared characteristics, namely library-based and region-based methods. We further conducted benchmark studies to evaluate the accuracy, sensitivity, coverage, and usability of NGS-based methods with molecular experimental datasets. Results show that BART, ChIP-Atlas, and Lisa have relatively better performance. Besides, we point out the limitations of NGS-based methods and explore potential directions for further improvement.

**Key points:**

- An introduction to available computational methods for predicting functional TRs from a query gene set.
- A detailed walk-through along with practical concerns and limitations.
- A systematic benchmark of NGS-based methods in terms of accuracy, sensitivity, coverage, and usability, using 570 TR perturbation-derived gene sets.
- NGS-based methods outperform motif-based methods. Among NGS methods, those utilizing larger databases and adopting region-centric approaches demonstrate favorable performance. BART, ChIP-Atlas, and Lisa are recommended as these methods have overall better performance in evaluated scenarios.

## Introduction

In the 1960s, foundational principles of gene transcriptional regulation were established through seminal work [1] and subsequent studies, which highlighted the pivotal role of transcriptional regulators (TRs) in controlling the transcription of hundreds or even thousands of genes [2, 3]. TRs are proteins encoded by genes and their regulation of transcription directly influences the expression levels of genes, playing a crucial role in various cellular processes including cell growth, differentiation, morphogenesis, and death [4–6]. A significant portion of these TRs are transcription factors (TFs), which bind to specific DNA sequences. The remainder of TRs consists of cofactors recruited by TFs and chromatin regulators that can alter the structure and function of chromatin [7, 8].

Evidence has linked multiple known diseases to the dysfunction of specific TRs [9, 10], highlighting the importance of identifying functional TRs in biological and medical research, which can aid the discovery of biomarker genes or potential therapeutic targets [11–13]. However, accurately determining the interactions between TRs and their targets requires significant computational and experimental efforts. Amid these challenges, one intriguing question that draws many researchers’ attention is how to accurately predict the TRs that regulate a user-provided gene set that can be derived from sources such as differentially expressed gene (DEG) analysis, gene ontology analysis, and pathway enrichment analysis.

In the past, a variety of computational approaches have been proposed to answer the aforementioned question based on microarrays and sequencing technologies. Among them, motif enrichment analysis was initially developed by associating over-represented transcription factor binding motifs (TFBMs) with cis-regulatory elements (e.g., promoters and enhancers), which regulate the transcription of nearby genes [14]. TFBMs are short DNA subsequences that have high affinities for specific TFs. If the TFBMs are over-represented, the related TFs are claimed to be functional regulators. Many methods have been developed following this rationale [15–17]. However, these methods share the following limitations: First, the high similarities of TFBMs among distinct TFs often make it challenging to decide which TFs actually bind the genomic loci [18]. Second, many TFs lack known binding motifs [19], which could not be analyzed by these published motif-based methods. Third, TFBMs are unable to capture the context-specific binding profile of a TF, which leads to a high false-positive rate in determining the actual binding sites in a particular cellular process. Therefore, motif enrichment analysis is not an ideal tool to search for TRs.

We further mention that the identification of TRs can serve as a downstream application of network construction analysis as well [20–22]. The realm of gene regulatory network reconstruction stands as a distinct research field, with comprehensive reviews providing deep insights into this domain [23–25]. Thus, these network construction methods were excluded from this review.

This review instead focuses on computational methods that can leverage recent advancement of next-generation sequencing (NGS) techniques, especially epigenomic techniques (e.g., ChIP-seq, ATAC-seq, and DNase-seq) to identify TRs that regulate a user-provided gene set. These methods are referred as NGS-based method in this review. Unlike motif-based methods, which screen the genome by computationally inferred motifs to find possible binding sites, using NGS data can provide evidence of actual binding sites within the genome.

Chromatin immunoprecipitation followed by sequencing (ChIP-seq) including TR ChIP-seq and Histone ChIP-seq is a widely accepted technique to examine genome-wide DNA-protein interactions. TR ChIP-seq datasets serve as the foundation of all NGS-based methods. A TR ChIP-seq dataset can accurately record the binding sites of one specific TR [26], and therefore makes it possible to establish the TR-gene connection.

Additionally, histone ChIP-seq techniques (e.g., H3K27ac ChIP-seq) and chromatin accessibility profiling techniques (e.g., DNase-seq, FAIRE-seq, and ATAC-seq) can measure genome-wide active cis-regulatory elements or accessible chromatin regions [27, 28]. Unlike TR ChIP-seq data required by all methods, data generated by these techniques cannot be directly associated with one specific TR, and so are only included by some methods for improving the prediction accuracy.

The interesting features extracted from these data are commonly termed as “peaks” [29], which are short epigenomic regions that show a significant enrichment of epigenetic marks or factors compared to the background levels. Peaks can indicate different features depending on the NGS techniques, such as TR binding sites (TR ChIP-seq), accessible chromatin regions (ATAC-seq, DNase-seq, FAIRE-seq), or sites of active promoters and enhancers (H3K27ac ChIP-seq).

Until now, over ten computational methods have been developed to predict TRs using NGS data. These methods are Cscan [30], ENCODE ChIP-seq significance tool[31], Enrichr [32], i-cisTarget [33], RegulatorTrail [34], BART [35], ChIP-Atlas [36], TFEA.ChIP [37], ChEA3 [38], MAGIC [39], and Lisa [40]. **Figure 1** illustrates the common workflow of these methods. The input is typically a query set of genes provided by a user. The background set comprises other genes in the same process that generates the query gene set (e.g., the query gene set consists of DEGs and the background set consists of non-DEGs). The reference database, composed of NGS data, functions as a unique ‘fingerprint’ repository for TRs and genes. Each TR ChIP-seq dataset in the database represents a distinct TR ‘fingerprint’. Users can compare a ‘fingerprint’ from a new cellular study against ‘fingerprints’ in this database and identify functional TRs based on their similarity. The methods used integrate input, background, and reference data to produce a TR ranking list, where higher ranks indicate a greater likelihood of the TR regulating the input gene set. In this sense, the methods discussed in this review mainly act as a ranking algorithm to rank TRs covered by their internal database based on the user input.

**Figure 1.**
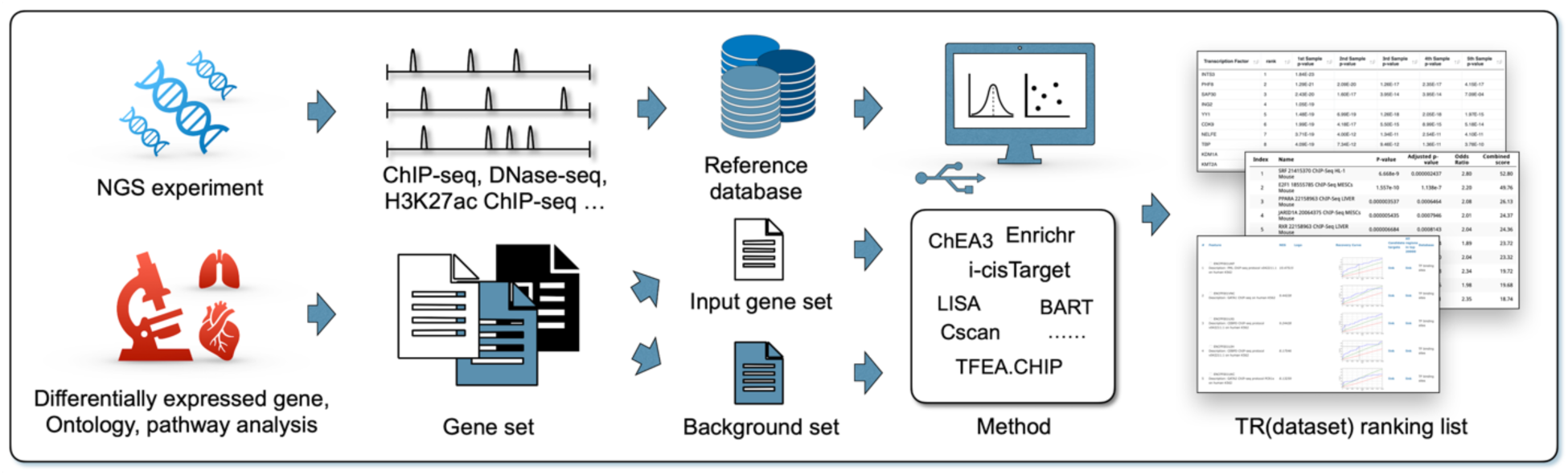
A general workflow of computational methods for predicting transcriptional regulators. Query gene set is produced by differentially expressed gene analysis, ontology, and pathway analysis. The reference database consists of NGS data from TR ChIP-seq, DNase-seq, and H3K27ac ChIP-seq. Eleven NGS-based computational methods so far have been developed to identify TRs. The final output is typically a TR ranking list.

Although each of these methods has been shown to perform well in certain applications, a unified and systematic evaluation has not been conducted yet. Considering their extensive use in thousands of publications so far [41–44], it is essential to elucidate their strengths and limitations by a benchmark study to avoid misinterpretation and inform the development of improved methods. The contribution of this review is three-fold. First, it offers a detailed overview of the NGS-based computational methods, highlighting their differences and similarities. Second, it provides an objective evaluation using an extensive collection of gene sets derived from TR perturbation experiments. Third, it systematically assesses key aspects of the methods, including accuracy, sensitivity, coverage, and usability, as well as limitations and directions for potential improvement.

## Categorization of NGS-based Methods

The NGS-based methods reviewed in this article fall into two main categories, each with two sub-categories: (i) library-based methods, relying on enrichment analysis of target gene (TG) libraries, subdivided into window-based and window-free approaches based on the main TG assignment algorithm, and (ii) region-based methods, which use cis-regulatory regions for TR prediction, further categorized into proximal-elements-centric and distal-elements-inclusive methods (**Figure 2**).

**Figure 2.**
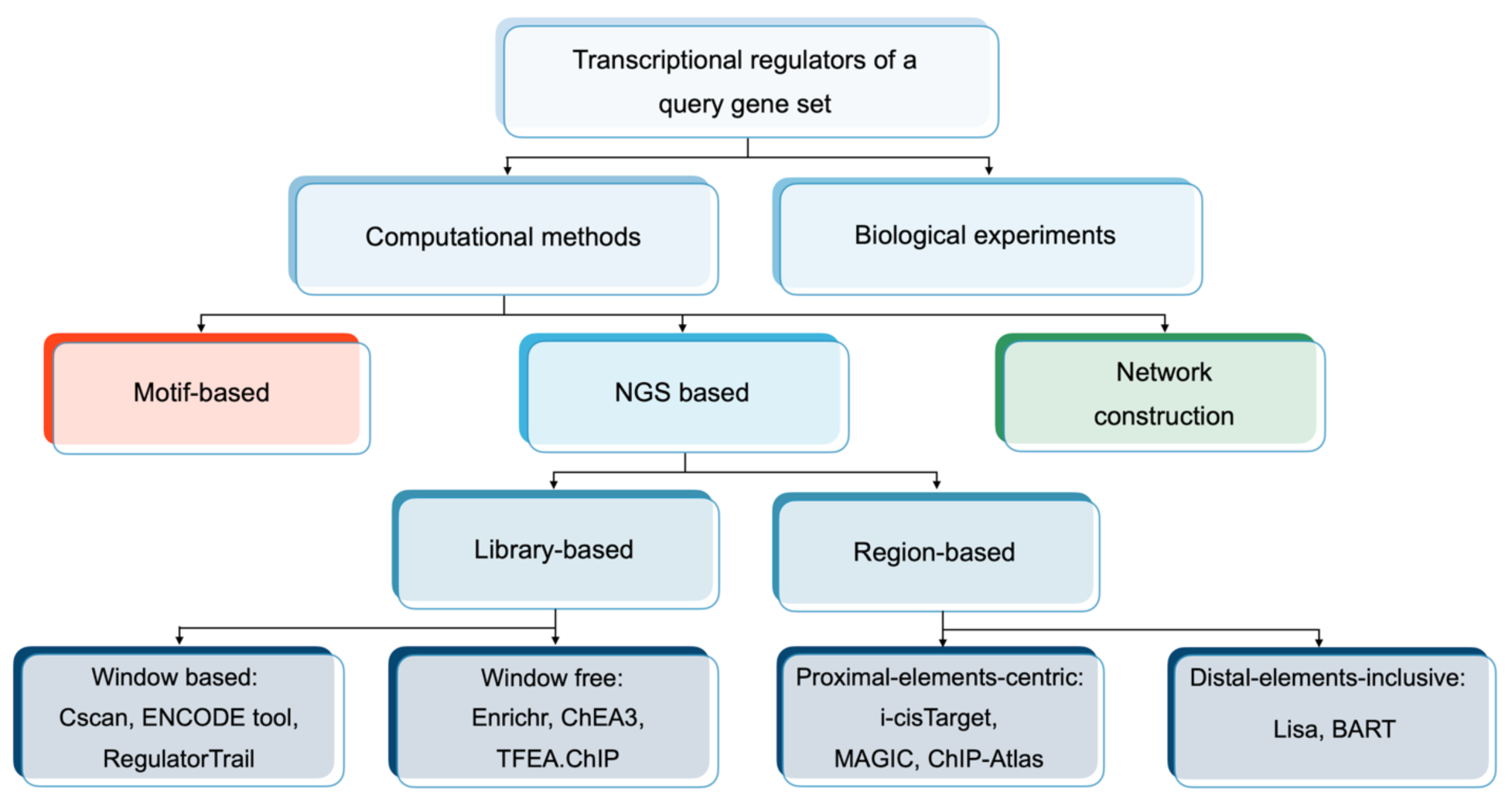
Categorization of computational methods for prediction of TRs mainly based on their approaches.

### TRs with multiple ChIP-seq datasets

Due to the uneven allocation of research resources across different TRs, while a TR ChIP-seq dataset is only linked to one specific TR, a single TR may be associated with multiple ChIP-seq datasets in an epigenomic database used. Methods including Cscan, Enrichr, ENCODE ChIP-seq significance tool (ENCODE tool), i-cisTarget, Lisa, and ChIP-Atlas generate a rank for each of the ChIP-seq datasets, even when they are linked to the same TR. Therefore, a final ranking list from any of these methods can contain multiple ranks for any TR with multiple ChIP-seq datasets. In contrast, the rest methods have an integration step to assign one unique rank to each TR. This integration can be achieved through various approaches, such as calculating the mean rank, selecting the highest rank, or employing statistical tests.

### Library-based methods

Predicting TRs that regulate a specific gene set requires establishing a link between TR binding sites and input gene symbols. Two main approaches exist: 1) converting TR binding sites to target gene sets followed by enrichment analyses to determine whether each target set is enriched by the input genes compared to the background set of genes, and 2) converting input gene symbols to their regulatory regions and identifying TRs with the highest proportion of binding peaks within these regions. Crucial to both strategies are the gene transcription start site (TSS), which serves as a reference point for mapping gene symbols to chromatin locations. This allows the calculation of the linear epigenomic distance between a TR binding peak and the TSS, a key measure for quantifying the TR-gene connection.

Library-based methods, including Cscan, ENCODE ChIP-seq significance tool, Enrichr, RegulatorTrail, ChEA3, and TFEA.ChIP, follow the first approach to create a TG library for TRs or TR datasets (**Figure 3**), where the library contains target gene sets identified from individual TR ChIP-seq experiments (Cscan, ENCODE tool, Enrichr, ChEA3) or integrated from multiple experiments for the same TR (RegulatorTrail, TFEA.ChIP). Genes in the input and background sets are dichotomized according to each target gene set in the library, and a statistical test, usually the Fisher exact test, is then applied to compute the statistical significance of each target gene set enriched with the input genes. Each TR or dataset is then ranked based on the statistical significance of their associated target gene set.

**Figure 3.**
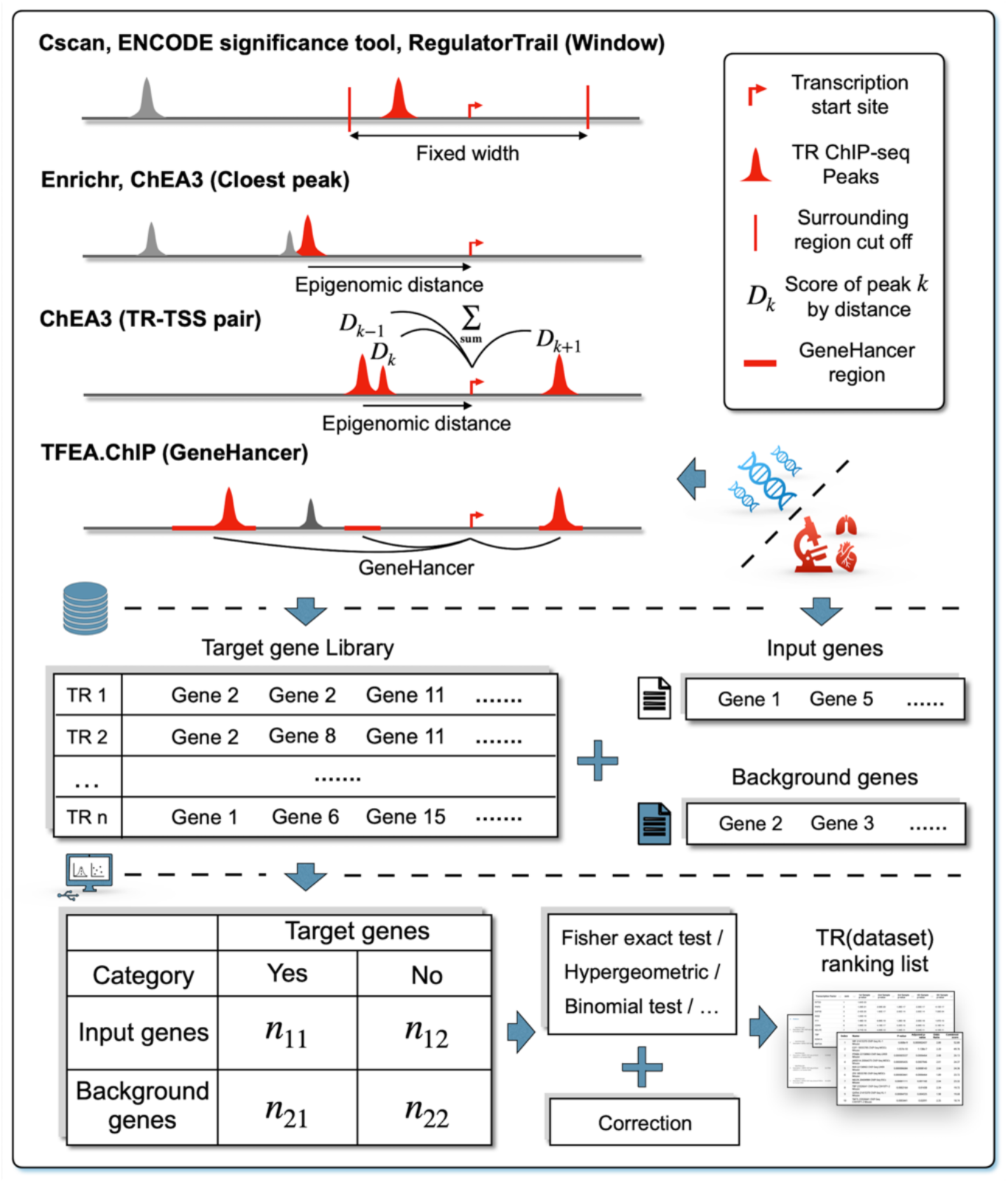
The common workflow of library-based methods. To set up a target gene library, TR ChIP-seq peaks are associated with the nearest genes, usually by measuring the genomic distance linearly to the transcription start site (TSS) or whether they fall into the surrounding region. The library can also be derived from experimental evidence. Fisher exact test is commonly used to decide whether the target gene set corresponding to each TR or dataset in the library is enriched with the input genes. The final output is a TR (dataset) ranking list based on the test significance.

Library-based methods mainly differ in their target gene assignment algorithms. Cscan, ENCODE tool, and RegulatorTrail, for example, employ a “window-based” approach. They define a fixed window (typically 50 bp to 10 kb) around each gene’s TSS. If a TR ChIP-seq dataset has peaks within this window for a particular gene, that gene is then classified as a target gene.

The other three methods mostly use a “window-free” approach. For example, Enrichr and ChEA3 focus on the proximity of peaks to genes. These methods sort peaks in a TR ChIP-seq dataset based on their linear epigenomic distances to the TSS of nearby genes and retain a certain number of genes with the closest peaks as targets. ChEA3 can further consider the density of peaks by employing a different method: first calculate a score for each TR-TSS pair by aggregating all the peaks of a TR ChIP-seq dataset within a specific range based on their distance to the TSS. Genes associated with TR-TSS pairs having the top 5% highest non-zero scores in each dataset are then designated as targets.

TFEA.ChIP is another “window-free” method, using the GeneHancer database that stores cis-regulatory elements of genes inferred from multiple data sources [45], such as FANTOM5 Ernagene expression correlation. These cis-regulatory elements are defined as “elite” regions, and TFEA.ChIP assigns the target genes of a TR based on whether any ChIP-seq dataset of the TR has peaks in these “elite” regions.

There are three key limitations to these six library-based methods. First, these methods use hard cutoffs to identify target genes. This binary approach may be overly simplistic. In Cscan, ENCODE ChIP-seq significance tool, and RegulatorTrail, the cutoff determines the window size, and any peaks outside the window are excluded from consideration. Similarly, Enrichr and ChEA3 use a cutoff to determine the number of genes retained in each ChIP-seq dataset. Since the selection of cutoff is subjective, different choices can result in different numbers of TGs and, subsequently, affect the prediction results.

Second, it is widely recognized that the 3D chromatin distance between an epigenomic peak and a gene promoter region, which can be measured by Hi-C or DNA-FISH [46], estimates their interaction more accurately than the one-dimensional linear distance between two loci (on a straight line) [47, 48]. However, most library-based methods use the linear distance to estimate the interactions, neglecting that the genome is organized in a complex three-dimensional structure, allowing distant locus to come in close physical contact [49, 50]. In addition, a significant fraction of binding sites has been shown to be neutral or non-functional in regulating gene expression even when located more proximal to a gene [51]. Thus, the dichotomization of genes based solely on the linear distance might lead to misleading conclusions.

Finally, library-based methods lack an approach to quantify the interaction between TRs and their target genes. Simply categorizing genes as targets and non-targets treats each TR-gene interaction as equal in importance. This neglects the fact that TRs can have different degrees of influence on gene expression. To fully understand how TRs regulate gene expression, researchers need methods that can not only identify target genes but also quantify the strength and functional significance of these interactions.

While the Fisher exact test is the primary statistical test used for enrichment analysis in librarybased methods, it is important to note some variations. Methods like RegulatorTrail and TFEA.ChIP offer alternative statistical tests such as the hypergeometric test, binomial test, or gene set enrichment analysis. Additionally, several methods (Cscan, RegulatorTrail, ChEA3, and TFEA.ChIP) incorporate FDR correction approaches like the Benjamin-Hochberg method to control false positives. Notably, Enrichr discussed the potential bias in the Fisher exact test due to varying gene set sizes and addressed it by employing z-scores that account for deviations from expected ranks, which are generated through simulations.

In summary, the performance of library-based methods heavily depends on the efficiency of TG assignment algorithms. However, there is no universally accepted algorithm. The enrichment analysis can further complicate this, with different statistical methods giving different ranking lists. In addition, the limitations mentioned above can severely hinder the performance of these librarybased methods.

### Region-based methods

The development of epigenomics now focuses on critical functions of cis-regulatory elements (CREs) [52], including promoters, enhancers, silencers, and insulators. These CREs can act as the binding sites for TRs and collaborate to regulate gene transcription [53], with promoters and enhancers being the two most important ones. Therefore, unlike library-based methods which convert the TR binding sites to target genes, region-based methods avoid using a TG assignment algorithm and instead focus on the enrichment of TR binding sites in regions that contain active CREs for the input genes and use additional information to quantify the TR-gene interaction.

Proximal-elements-centric methods, including i-cisTarget, MAGIC, and ChIP-Atlas, mainly use proximal CREs, which are essential for basic gene transcription, to predict TRs (**Figure 4**). Proximal CREs are regulatory sequences found close to the gene they regulate. Promoters, for instance, are a type of proximal CRE located immediately upstream of a gene’s TSS, which can serve as the “on-off” switch to initiate gene transcription [54]. Due to their proximity to the genes that they regulate, proximal CREs often have a more direct and immediate impact on gene expression compared to distal regulatory elements located farther away from the gene locus. Past evidence suggests TRs bound to the proximal CREs usually regulate the nearby genes [55]. Therefore, the rationale behind these methods is that functional TRs should have more peaks or high-signal peaks that match the proximal CREs of their regulated genes.

**Figure 4.**
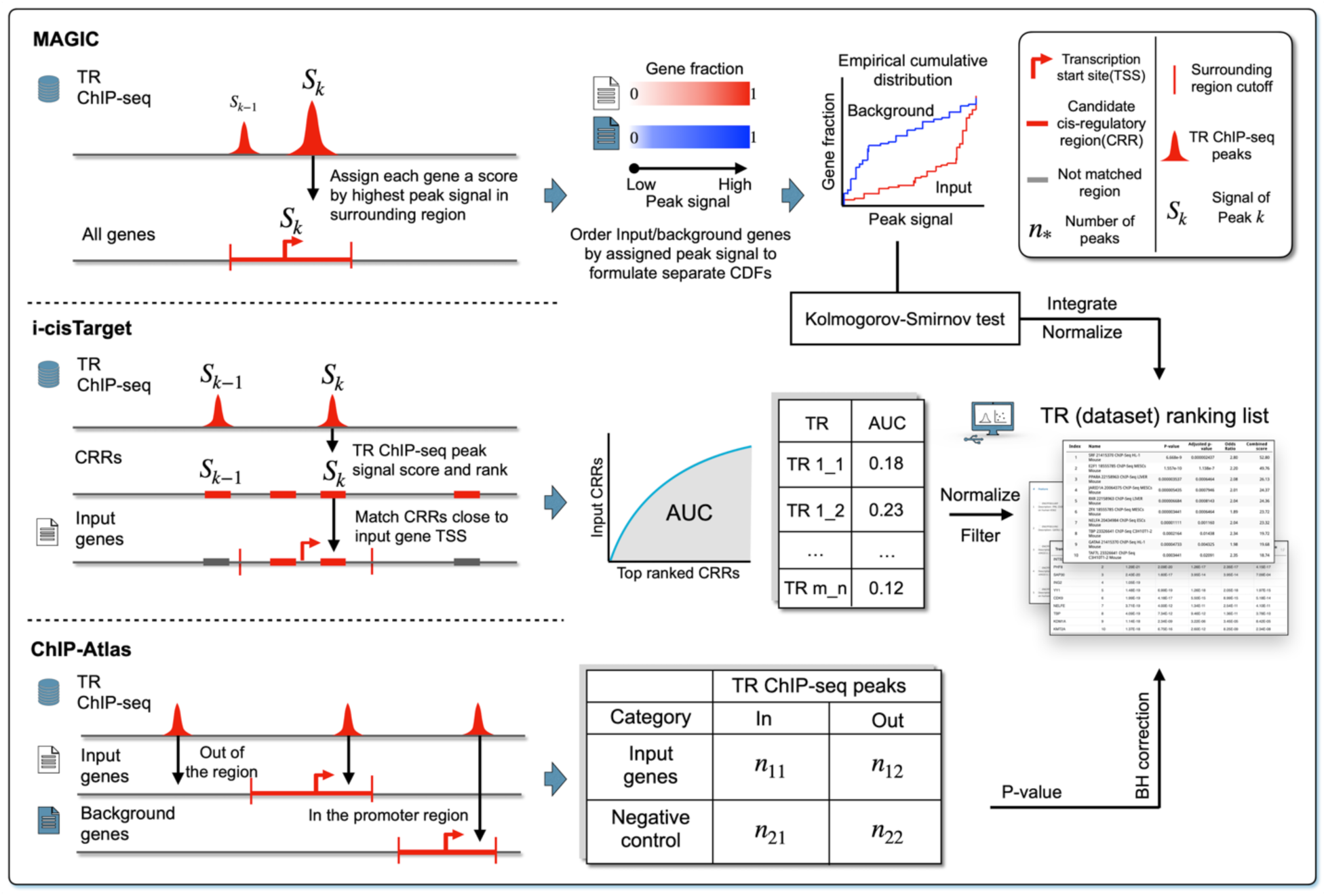
Workflow of the proximal-elements-centric method. All three methods focus on the regions that surround the gene body or gene’s TSS. TR ChIP-seq datasets with more peaks enriched in these regions would be ranked higher.

i-cisTarget and MAGIC focus on peaks with high signal values. The peak signal is calculated based on the number of sequenced reads aligned to the peak position; a high signal value suggests a strong TR-DNA interaction. i-cisTarget first predefined many candidate cis-regulatory regions (CRRs) that may contain CREs based on public sources, mainly DNaseI hypersensitive sites (DHSs). i-cisTarget scores and ranks these CRRs for each TR ChIP-seq dataset based on peak signals within the regions. Then, it calculates an area under the curve (AUC) value for each dataset by counting how many top-ranked CRRs can match the CRRs around the TSSs of input genes. Raw AUCs for the TR ChIP-seq datasets are filtered according to a threshold value and then normalized to obtain final scores that yield the final ranking list of these datasets.

MAGIC assigns each gene in the input and background the highest signal value among peaks that fall into its surrounding region, and the step is repeated for each TR ChIP-seq dataset. Next, the input and background genes are ranked based on these values to create two separate empirical cumulative distribution functions (eCDFs) for each dataset. Kolmogorov-Smirnov test is used to identify the TR ChIP-seq datasets that most effectively differentiate between the two eCDFs.

ChIP-Atlas does not use signal values. Instead, it aims to identify which TR ChIP-seq dataset has a higher proportion of peaks enriched in the promoter regions of the input genes than in those of the background genes. Even though a one-tailed test is more appropriate here, ChIP-Atlas applies a two-tailed Fisher exact test to determine the statistical significance for each TR ChIP-seq dataset, which is used to obtain the final ranking list.

A significant limitation of these methods lies in the unreliability of using signal strength to measure functionality. While strong peak signals often correlate with stronger binding affinity or functionality [56], they aren’t a perfect measure. This is because ChIP-seq peak signals are computed based on an average across millions of cells. Peaks with low or medium signal could represent highaffinity binding sites in a specific subpopulation of cells, while high signal peaks might reflect only moderate affinity sites present in all cells [57]. Furthermore, emerging evidence suggests that even low-affinity binding sites can play crucial roles in regulating gene transcription [58]. Therefore, a strong signal alone does not guarantee a functionally important regulatory element.

In addition to proximal CREs, distal regulatory elements like enhancers also can play crucial roles in gene transcription regulation [59, 60]. Located tens of thousands of base pairs away from their target genes, these distal CREs modulate the rate and timing of gene expression, ensuring precise control. Distal CREs are highly variable and more sensitive to environmental factors compared to proximal CREs. While incorporating distal CREs into TR prediction models could improve accuracy, directly measuring the long-range interactions between these elements and their targets remains challenging due to limited data availability.

The distal-elements-inclusive methods, including BART and Lisa, attempt to incorporate the activity of distal CREs by using H3K27ac ChIP-seq datasets (**Figure 5**). H3K27ac is a type of Histone protein used to mark active enhancers and promoters [27]. Instead of directly identifying which TR has more peaks enriched in the proximal regions, the two methods first select a few H3K27ac ChIP-seq datasets that can be predictive for the transcription of input genes, which serve as genome-wide CREs activity landscape. The rationale of the two methods is that a functional TR should have a higher number of peaks aligned with the high signal peaks in these selected H3K27ac ChIP-seq samples.

**Figure 5.**
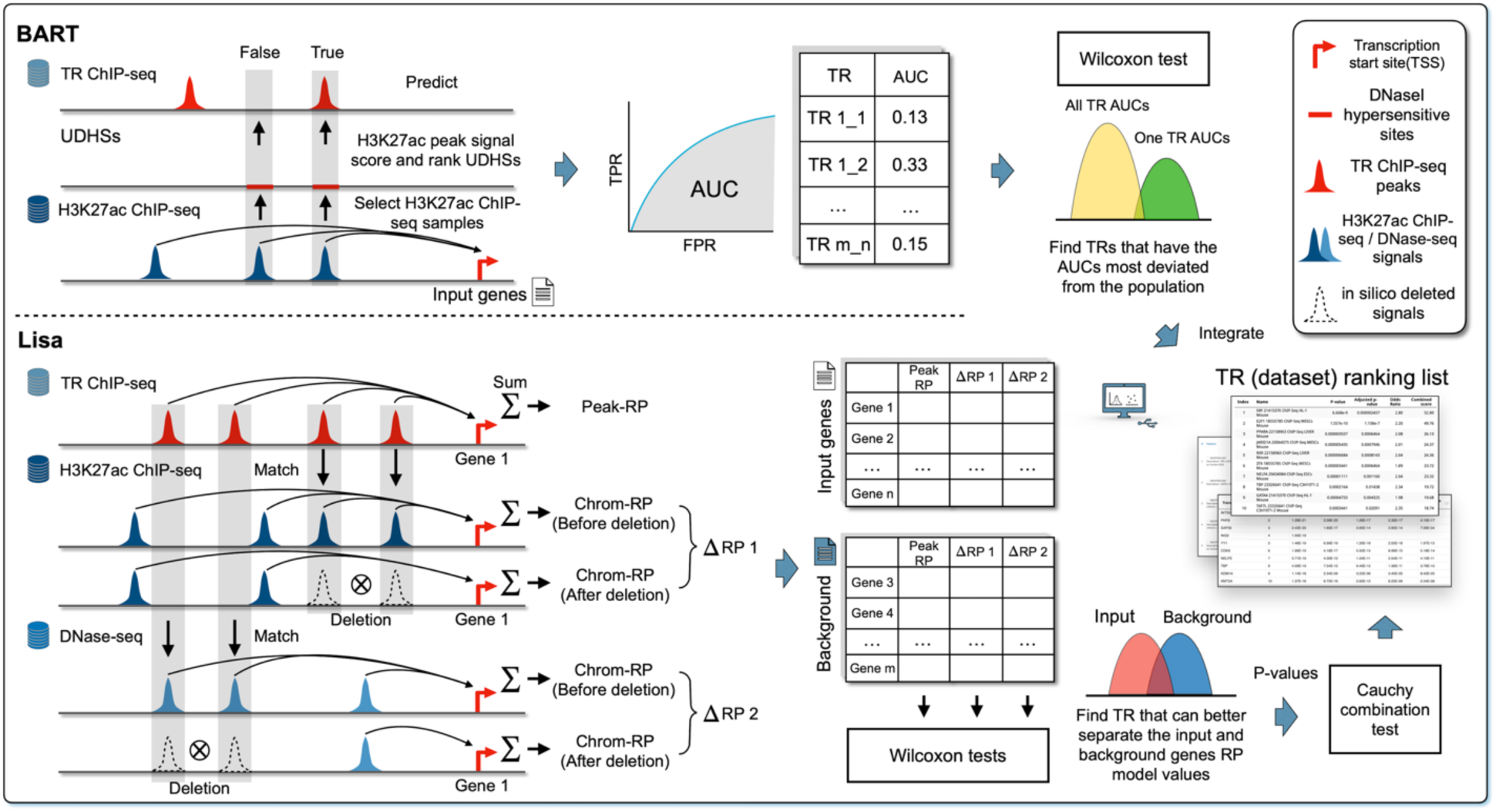
The core idea of BART and Lisa. Distal-elements inclusive methods first select predictive H3K27ac ChIP-seq (or DNase-seq) samples that quantify active distal cis-regulatory elements, by summarizing the H3K27ac ChIP-seq (or DNase-seq) peak signals around a gene’s TSS. Next, statistical methods are applied to examine which TRs align more frequently with the high signal peaks from these selected samples.

In the selection step, a regulatory potential (RP) model is implemented in both methods, which counts the H3K27ac signals within the surrounding region of each gene to assign an RP score. Therefore, each H3K27ac dataset has a unique RP score for each individual gene. Following this, predictive H3K27ac datasets are chosen using stepwise logistic regression. That is, the two methods model the presence of a gene in the input list as a binary outcome (yes/no). The gene’s RP scores across all H3K27ac datasets are used as potential predictors. Then, using a stepwise selection method, the two methods choose a subset of these datasets that best explains whether the gene is included in the input list.

After selecting predictive H3K27ac datasets, BART collects genome-wide DHSs and for each DHS, computes a score by the weighted combination of signals from the selected datasets, and then uses these scores to predict the peak presence in each TR ChIP-seq dataset and calculates the AUC value using false positive rate (FPR) and true positive rate (TPR). The AUC values of the ChIP-seq datasets are grouped by TR and each TR’s AUC values are compared to the distribution of AUC values from all TRs through the Wilcoxon rank sum test. Finally, BART assigns each TR a unique rank by averaging the ranks of the maximum AUC value, the p-value from the test, and the standardized Wilcoxon statistic score that is derived using TR-specific mean and standard deviation calculated by running BART on more than 500 pre-collected gene sets.

In addition to H3K27ac datasets, Lisa selects predictive DNase-seq datasets as well; RP scores are referred to as “chrom-RP” for the selected datasets of either type. After the selection, for each type, Lisa matches the peaks from each TR ChIP-seq dataset with those from the selected datasets, deleting all matched signals to reproduce the chrom-RP for each gene. Then the fitted logistic regression model is used to calculate the chrom-RP change before and after the deletion, denoted as ΔRP for each gene. Lisa further calculates a “Peak-RP” for each gene by using peaks from the TR ChIP-seq dataset. Therefore, each TR ChIP-seq dataset has two ΔRPs, one from selected H3K27ac samples and the other from selected DNase-seq samples, and one peak-RP for each gene in the input and background sets. Finally, for each TR ChIP-seq dataset, ΔRPs and peak-RPs are analyzed by three separate Wilcoxon tests to test whether there is a significant difference between the input and background, and the p-values are integrated by Cauchy combination test, the significance of which is used to derive a final ranking list of all TR ChIP-seq datasets.

While both BART and Lisa integrate data from other NGS techniques to analyze distal CREs, they still prioritize CREs closer to genes. Also, the presence of the H3K27ac histone mark alone might not reliably indicate CRE activity [61]. This is similar to the limitations of using peak signals from TR ChIP-seq, where a strong signal cannot guarantee functional binding. Consequently, accurately quantifying CRE activity or functionality of TR binding sites to their target genes remains a significant challenge.

Finally, the uneven allocation of research resources across different TRs also leads to a bias where a select few TRs and cell lines have been studied in much greater depth than others. This favoritism towards certain TRs (like CTCF and TP53) and cell lines (like HeLa and K562) often stems from research preferences or the availability of experimental materials. Since these methods rely on multiple statistical tests to assign significance to each dataset, they tend to favor TRs with extensive data coverage over those with fewer datasets.

**Table 1** provides a summary of the eleven NGS-based methods examined in this study. The majority rely on data from the ENCODE [62], a trusted source for high-quality NGS data. Methods like LISA and ChIP-Atlas expand their scope by incorporating NGS datasets from individual studies. This approach offers wider TR and cell line coverage but may introduce potential variability in data quality. The evolution of these methods reflects a clear trajectory. Initially librarybased, they shifted to region-based, and now employ increasingly sophisticated strategies. This highlights the field’s continuous adaptation to new NGS technologies and its ability to address increasingly complex scientific questions.

**Table 1.** The summary of evaluated eleven NGS-based methods.

| Method (Year) | NGS data source | Implementation | Major method to associate TFs with genes | Primary statistical method | Method categorization |
| --- | --- | --- | --- | --- | --- |
| Cscan (2013) | ENCODE | Web tool | Peaks fall into the surrounding fixed width window of gene TSS | Fisher exact test | Library-based |
| ENCODE ChIP-seq significance tool (2013) | ENCODE | NA | Peaks fall into the surrounding fixed width window of gene TSS | Hypergeometric test | Library-based |
| Enrichr (2013) | ENCODE / ReMap / Literature | Web tool | Epigenomic distance between peaks and gene TSS | Fisher exact test / Corrected FET | Library-based |
| i-cisTarget (2015) | ENCODE | Web tool | Peak signal to score pre-defined CRRs | Recovery analysis | Region-based |
| RegulatorTrail (2017) | Multiple sources | Web tool | Peaks fall into the surrounding fixed width window of gene TSS | FET / Binomial test / Hypergeometric test | Library-based |
| BART (2018) | CistromeDB | Web tool / Python | RP model to score pre-collected DHSs | Wilcoxon rank sum test | Region-based |
| ChIP-Atlas (2018) | NCBI / DDBJ / ENA | Web tool | Proportion of peaks fall into gene promoter region | Fisher exact test | Region-based |
| TFEA.ChIP (2019) | ENCODE / ReMap | Web tool / R | Peaks fall into the "elite" regions from GeneHancer | FET / Kolmogorov Smirnov test / GSEA | Library-based |
| ChEA3 (2019) | ENCODE / ReMap / Literature | Web tool | Epigenomic distance between peaks and gene TSS | Corrected Fisher exact test | Library-based |
| Lisa (2020) | CistromeDB | Web tool / Python | RP model with in-silico deletion method | Wilcoxon rank sum test / Cauchy combination test | Region-based |
| MAGIC (2020) | ENCODE | Python | Peak signal to rank the genes | Kolmogorov-Smirnov test | Region-based |

## Methods

### Benchmarking study design

We systematically compared the performance of the NGS-based methods reviewed in the previous section under a unified evaluation framework encompassing four key dimensions: accuracy, sensitivity, coverage, and usability. Also included are three motif-based methods, Pscan [15], HOMER [17], and RcisTarget [63] that can accept a gene set as input and prioritize potential TRs through motif enrichment analysis. These methods use motif libraries of known TRs and assign a higher rank to a TR whose associated motifs are determined to be over-represented. By including motif-based methods, it is possible to assess whether using NGS data can indeed enhance the prediction accuracy.

We used gene sets sourced from KnockTF, a comprehensive human gene expression profile database generated from TR knockdown/knockout experiments [64]. Differentially expressed genes derived from perturbation experiments often serve as a benchmark for evaluating the performance of TR-related computational methods [65]. In the database, each TR perturbation experiment has at least two control samples and two treatment samples, where Limma [66] was used to derive the differentially expressed (DE) gene set. In total, we retrieved 570 DE gene sets from the database, covering 308 known TRs. We further selected the top 200, 600, and 1000 most significant genes from each input gene set, respectively. The top 200 genes are used to evaluate accuracy as these genes should be highly correlated with the perturbed TR. The other two are used to evaluate sensitivity of the methods against noisy input. We transformed the annotated genes to the required format following the manual or instructions. For each method considered, we used the default settings when applicable, otherwise the parameters were chosen according to the documentation.

To assess a method’s accuracy, we focused on its ability to rank known, perturbed TRs at the top of its generated ranking list. This aligns with the user’s primary interest: identifying the most relevant TRs at the forefront of the results. We first selected a range of threshold values (*K*) for “top *K* positions” (*K*=10, 50, 100). For each *K* value, we then applied four widely recognized metrics designed for evaluating ranking algorithms to assess the ranking results from the different methods [67]. Given gene lists *i* = 1,2, …, N, N is the total number of evaluated gene lists. The terms ranked above threshold are ordered as *k* = 1,2, …, *K* by their ranks. The relevance of each term in the ranking list is then denoted as *Ri_k_* where *Ri_k_* = 1 if the term is indeed the perturbed TR, and 0 otherwise. The total number of relevant terms in the ranking list *i* is denoted as *ni_k_*, where 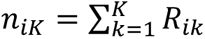 does not exceed the number of TR ChIP-seq data sets associated with the perturbed TR. The metrics used in our evaluation include (1) Hit rate at *K*, which calculates the proportion of ranking lists that contain at least one relevant term above the threshold,

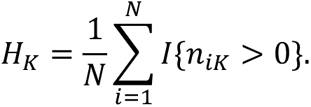

(2) Mean reciprocal rank (MRR) at *K*, which values the first relevant term in each ranking list,

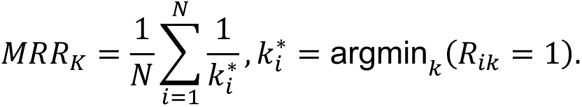

(3) Mean average precision (MAP) at *K*, which measures a method’s capability of ranking relevant terms at the top, penalizing deviations. For a ranking list *i*, the metric first calculates the precision *P_i_* (*k*) at every *k*. It then sums over all *k*, further divided by *ni_k_* for list *i*. Finally, it averages across all the ranking lists to get the overall precision.

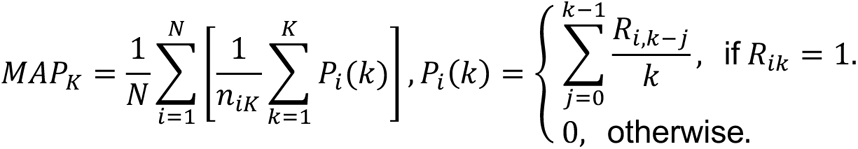

(4) Mean normalized discounted cumulative gain (mNDCG) at *K*, which also considers the position of relevant terms in a ranking list while giving more weights to the terms placed higher.

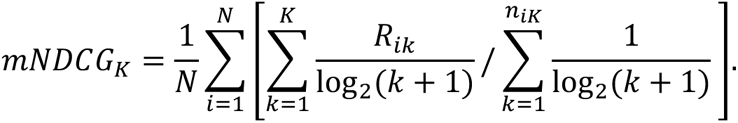

Each of the four metrics has a different focus. The hit rate measures in general whether a method can identify the perturbed TRs. While MRR, MAP and mNDCG consider the relevant positions of perturbed TRs in the top, MRR focuses exclusively on the ranking of the first relevant term. In case a method ranks multiple datasets linked to the same TR above the threshold, MAP and mNDCG evaluate the rankings of all relevant terms but assign different weights and penalties. In addition to the ranking quality metrics that consider the absolute positions of perturbed TRs, we also counted the number of ranking lists that have perturbed TRs shown in the top 1%, top 5%, and top 10%. The accuracy of each method is ranked by their average performance over all metrics and thresholds.

We also evaluated the sensitivity of methods to noisy input, focusing on their ability to produce consistent and reliable results. We tested methods with *G* = 200, 600, 1000 genes that are most significant, respectively. The size of the input gene set is the major parameter that can be controlled by users. As *G* increases, the number of non-significant genes in the gene set tends to increase, which may negatively affect the performance. In addition, we investigated whether TRs with multiple ChIP-seq datasets are significantly ranked higher compared to TRs with only one dataset by using Wilcoxon test. The p-values derived from same size gene set are integrated by Stouffer’s Z-score method.

Beyond accuracy and sensitivity, a method’s coverage is crucial. This refers to the breadth of TRs and cell lines represented in its internal NGS database. Without relevant data, a method cannot rank a specific TR, excluding it from results. Therefore, coverage directly impacts performance – broader coverage allows for wider analysis and a greater chance of identifying relevant TRs. Methods with continually updated databases tend to have better coverage, reflecting the latest discoveries.

It is worth noting that even high-performing methods may face practical implementation challenges. Therefore, usability is critical and depends on ease of implementation, quality of documentation, and code clarity. To assess usability, we defined nine criteria, each scored as 0 (not met) or 1 (met). These criteria address key user considerations when selecting a method: To assess usability, we defined nine criteria, each scored as 0 (not met) or 1 (met). These criteria address key user considerations when selecting a method: (1) a detailed manual or tutorial providing clear instructions, (2) illustrative examples helping users understand the analysis workflow, (3) availability of a CLI/API or R/Python package facilitating large-scale analyses, (4) a user-friendly web tool allowing direct method execution, (5) open-source code allowing for customization and transparency, (6) active maintenance ensuring the method remains functional, (7) visualization enhancing user comprehension of analysis results, (8) data import accepting gene symbols without requiring complex conversions, and (9) minimal unexpected errors during use ensuring a smooth user experience.

Within this evaluation framework, we benchmarked thirteen methods. Note that we excluded the ENCODE ChIP-seq significance tool due to being outdated and inaccessible. For Enrichr, we used the highest ranks from four libraries. For ChEA3, we followed the original publication’s recommended approach of utilizing the integrated mean rank. Additionally, it’s important to note that i-cisTarget web tool employs a filtering step, therefore only datasets that pass this filter were included in our evaluation.

## Results

### Accuracy

Figure 6 provides an overview of the performance of thirteen methods evaluated for ranking perturbed TRs. In summary, no single method stood out as the best in all evaluation metrics. However, actively updated or recently developed methods were observed to perform better, perhaps due to their larger accumulation of NGS data and more sophisticated algorithm design. Some general trends and potential issues were also observed, as detailed below.

**Figure 6.**
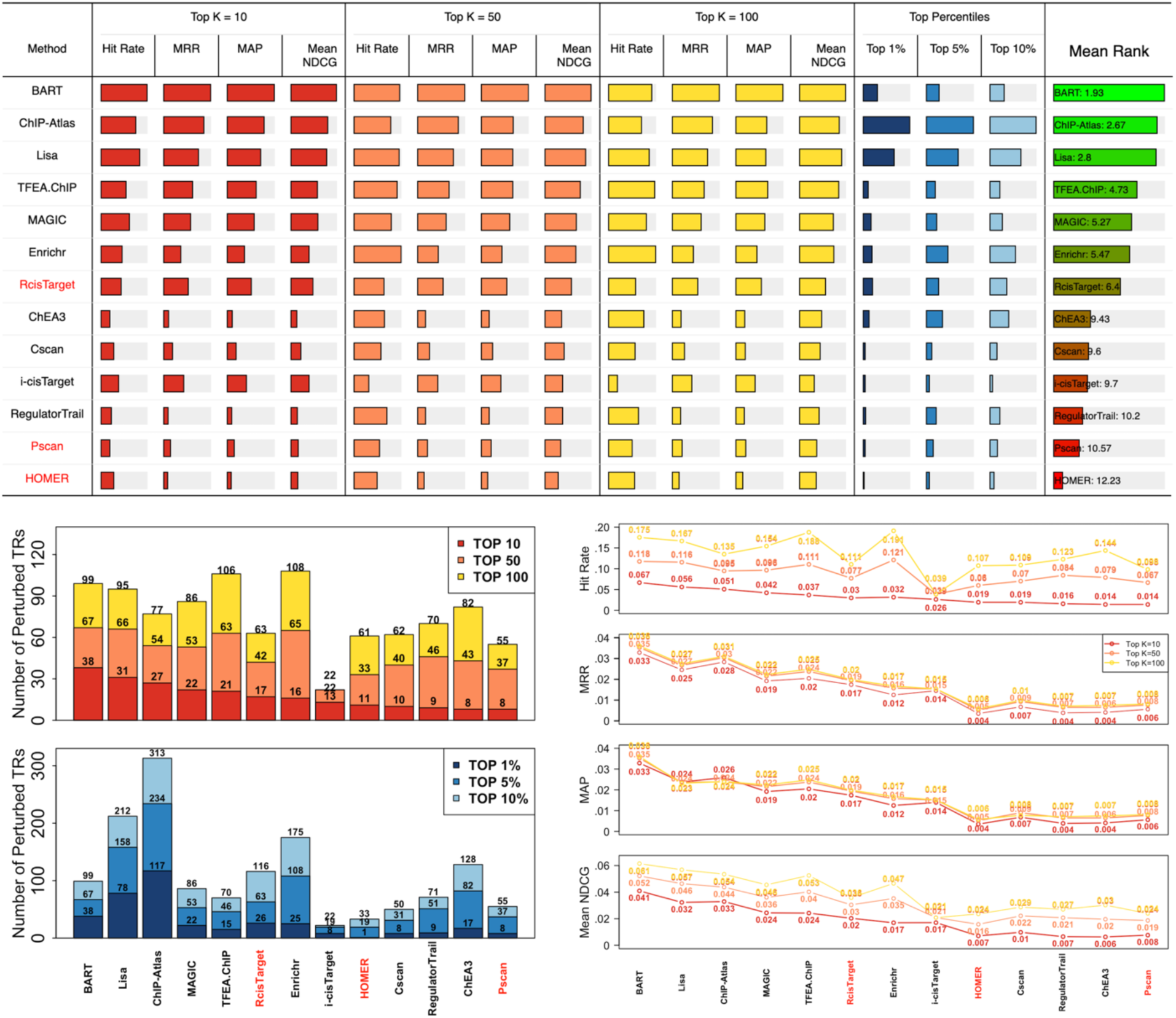
Comparison of accuracy. Methods with red labels are motif-based. BART, ChIP-Atlas, and Lisa demonstrate the best overall performance across different scenarios. The NGS-based methods generally outperform the motif-based methods. All methods ranked fewer than 10% of perturbed TRs within the top 10 positions, suggesting a room for further improvement.

First, BART, ChIP-Atlas, and Lisa are recommended in general as they are the top three performing methods in most scenarios. For example, BART, ChIP-Atlas, and Lisa have higher hit rate, MRR, MAP, and NDCG values for three selected thresholds. They also performed well by considering the top percentile rankings. The good performance is partially credited to the extensive collections of NGS data: ChIP-Atlas contains over 30,000 human TR ChIP-seq datasets, BART and Lisa have around 7,000 human datasets, while other methods only have 1,000 to 3,000 datasets. Furthermore, the region-based approach might also play a pivotal role in enhancing their accuracy. TFEA.ChIP, MAGIC, Enrichr, and RcisTarget also show good performance under certain metrics. Besides, NGS-based methods outperform motif-based methods on average. No motif-based method evaluated can surpass the performance of the top-performing NGS-based methods (e.g., BART, ChIP-Atlas, and Lisa), regardless of the metrics and thresholds used.

Combining results from various methods is believed to be more effective than relying on a single method due to differences in databases and algorithms used. We next explore combining two methods to determine the most optimal results. Figure 7 illustrates the number of perturbed TRs that are ranked in the top 10, 50, and 100 and the top 1%, 5%, and 10% in at least one TR ranking list when two methods are used (200 input genes used; see **Figure S1** and **Figure S2** for 600 and 1000 input genes). We noticed that Lisa, TFEA.ChIP, BART, and ChIP-Atlas can be combined to capture more perturbed TRs. In most cases, combining any of the two methods might yield better performance than using individual methods alone.

**Figure 7.**
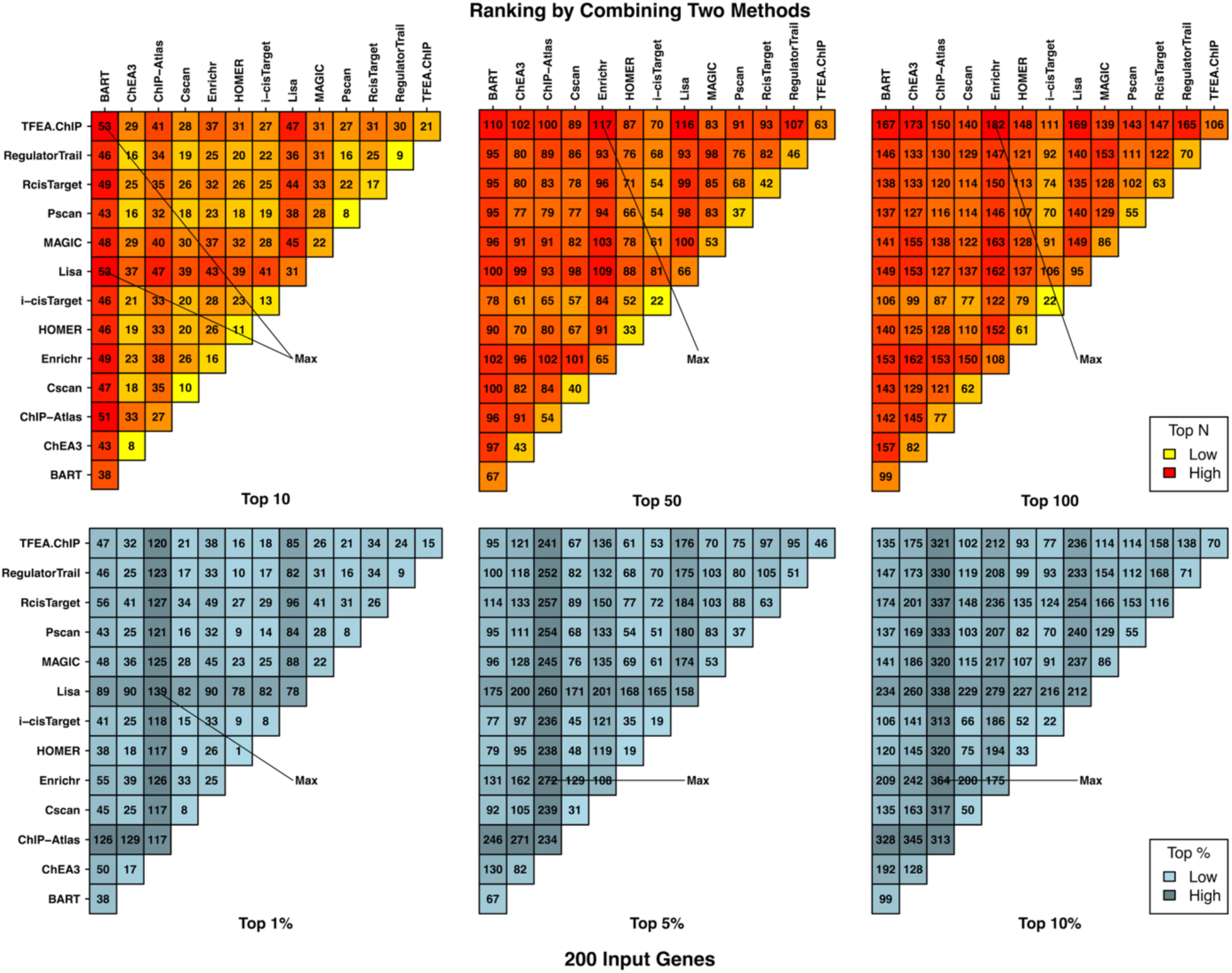
The number of perturbed TRs ranked to top positions by either method in the combination of two methods (200 input genes used). The combination of any two from BART, ChIP-Atlas, Lisa, Enrichr, and TFEA.ChIP can achieve the best or close to the best performance.

Figure 8 reveals a tendency among many methods to consistently rank certain TRs highly, regardless of the input gene set. However, these “favorite” TRs are rarely the experimentally perturbed TRs we want to identify. Prioritizing irrelevant TRs can overshadow true functional regulators, pushing them lower in the ranking list. This issue is most pronounced in RegulatorTrail, Cscan, and MAGIC. Thus, even if top-ranked TRs seem expected or familiar, we recommend users cross-check them against a method’s “favorite” TR list before proceeding with further analysis.

**Figure 8.**
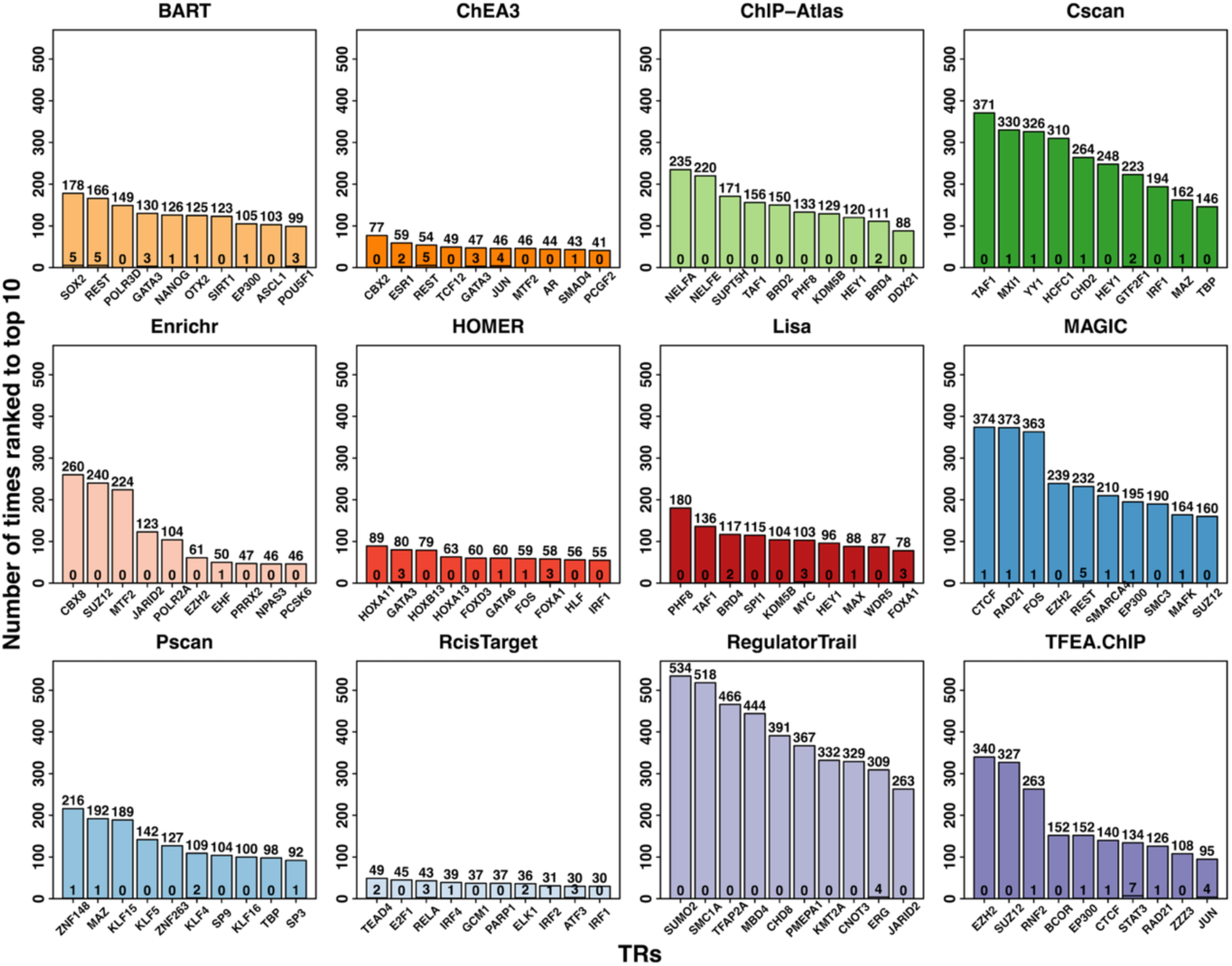
TRs that are most frequently ranked to the top 10 positions by different methods. The number at the top of each bar indicates how many times the corresponding TR is ranked in the top 10 and that at the bottom indicates how many times that it is indeed the perturbed TR (out of 570). i-cisTarget was excluded as the number of TRs in the final ranking list is often less than 10 due to the threshold.

### Sensitivity

Figure 9 presents results from the sensitivity analysis with three input gene set sizes. As the input size increases, most methods have reduced performance as we expect, indicating that the inclusion of more genes into the input set will lead to decreased performance of most methods. Therefore, it should be cautious when selecting genes used as input for these methods. The number of input genes itself can be a factor affecting the method performance. It is noted that HOMER, Pscan, and ChEA3 show relatively stable performance compared to others. However, given that accuracy remains the major consideration, among the top-performing methods (e.g., Lisa, ChIP-Atlas, and BART), Lisa is less sensitive to the change of input size.

**Figure 9.**
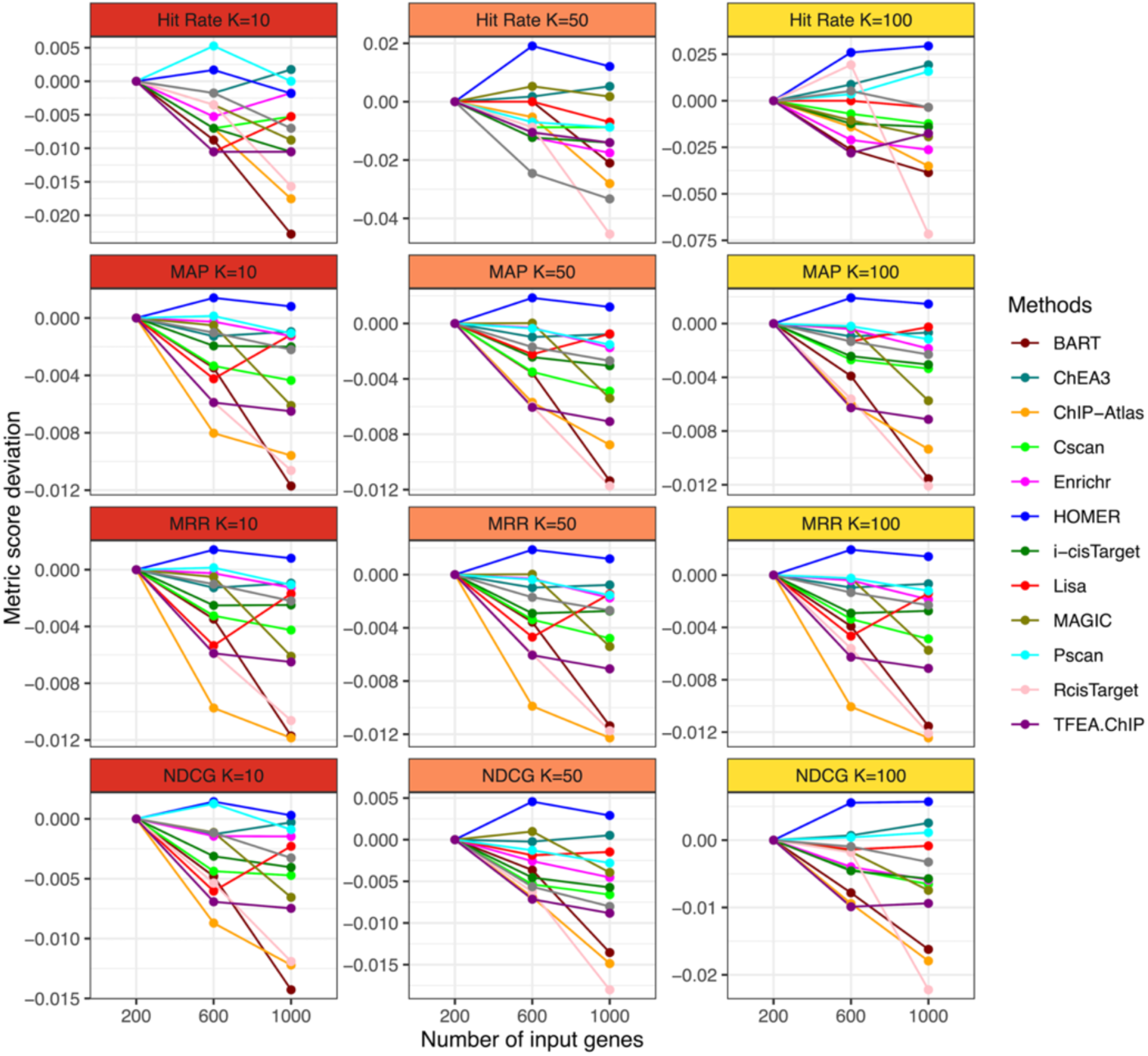
As the size of the input gene set increases, the majority of methods demonstrate a decline in performance. Notably, HOMER, Pscan and ChEA3 show relatively stable or even better performance with increased number of input genes. For the top 3 performing methods in the evaluation of accuracy, Lisa displays the most consistent performance compared to BART and ChIP-Atlas.

Except for the input gene set size, another major concern is the uneven allocation of research resources, as some TRs have been extensively investigated (**Figure S3**). We selected 20 TRs with the largest numbers of ChIP-seq datasets and 20 TRs associated with just one dataset randomly from the ENCODE database. For each gene set, we employed a one-sided Wilcoxon rank-sum test to determine if the 20 TRs with multiple datasets tend to be ranked higher than those with a single dataset. Additionally, we excluded gene sets that feature selected TRs as the perturbation targets, ensuring that each gene set could be considered a randomly generated set that is irrelevant to any of the selected TRs. The findings, detailed in **Table S1**, reveal that most methods indeed tend to favor TRs associated with a larger number of datasets. ChEA3 is the only method that does not give significant results, which might be due to the integration step of using mean ranks of all libraries as the final rank. This bias might explain why certain TRs consistently appear at the top as shown in Figure 8.

Finally, we illustrated how the value of a cutoff can affect the rankings of individual perturbed TRs that are expected to be ranked high by any appropriate methods (**Figure S4 – S5**). We sourced ∼2000 TR ChIP-seq datasets from the ENCODE project and implemented similar workflows as Cscan and Enrichr, both library-based methods, to identify target genes, followed by a one-sided Fisher exact test for each TR to produce the rank based on its significance. To assign target genes, Cscan uses windows of a predetermined width and Enrichr retains a prefixed number of genes based on the genomic distance, as discussed before. The figures show that, no matter which quantity is used as the cutoff, the individual ranks of some perturbed TRs can change drastically as we vary the cutoff, which again confirms that the use of a hard cutoff may pose a challenge in these methods.

### Coverage

With the detailed analysis of accuracy, it is evident that coverage plays a crucial role in the effectiveness of methods. The extent to which these methods can incorporate up-to-date NGS data can be a significant factor contributing to the performance (Figure 10). First, Figure 10 **A** and **B** show that the publication time has a positive correlation with the ranking performance and TR coverage, which confirms that the continuing efforts from generations of researchers have improved the prediction performance of algorithms to address the problem. Second, Figure 10 **C** displays the number of perturbed TRs that are missing in the TR ranking list versus the publication date for each method. As we expect, more recently developed methods generally have lower numbers of missing TRs. Figure 10 **D** plots the number of unique terms displayed in the TR ranking list of each method. Considering that the benchmark gene sets used in this study only cover a limited number of known TRs, a high number of missing TRs coupled with a low number of available terms might suggest an insufficient coverage of TRs. Notably, ChIP-Atlas, Lisa, and BART demonstrated their relatively comprehensive coverage of NGS data, which resonates with their better accuracy observed before, indicating the method performance is positively correlated with the number of covered TRs in method’s database.

**Figure 10.**
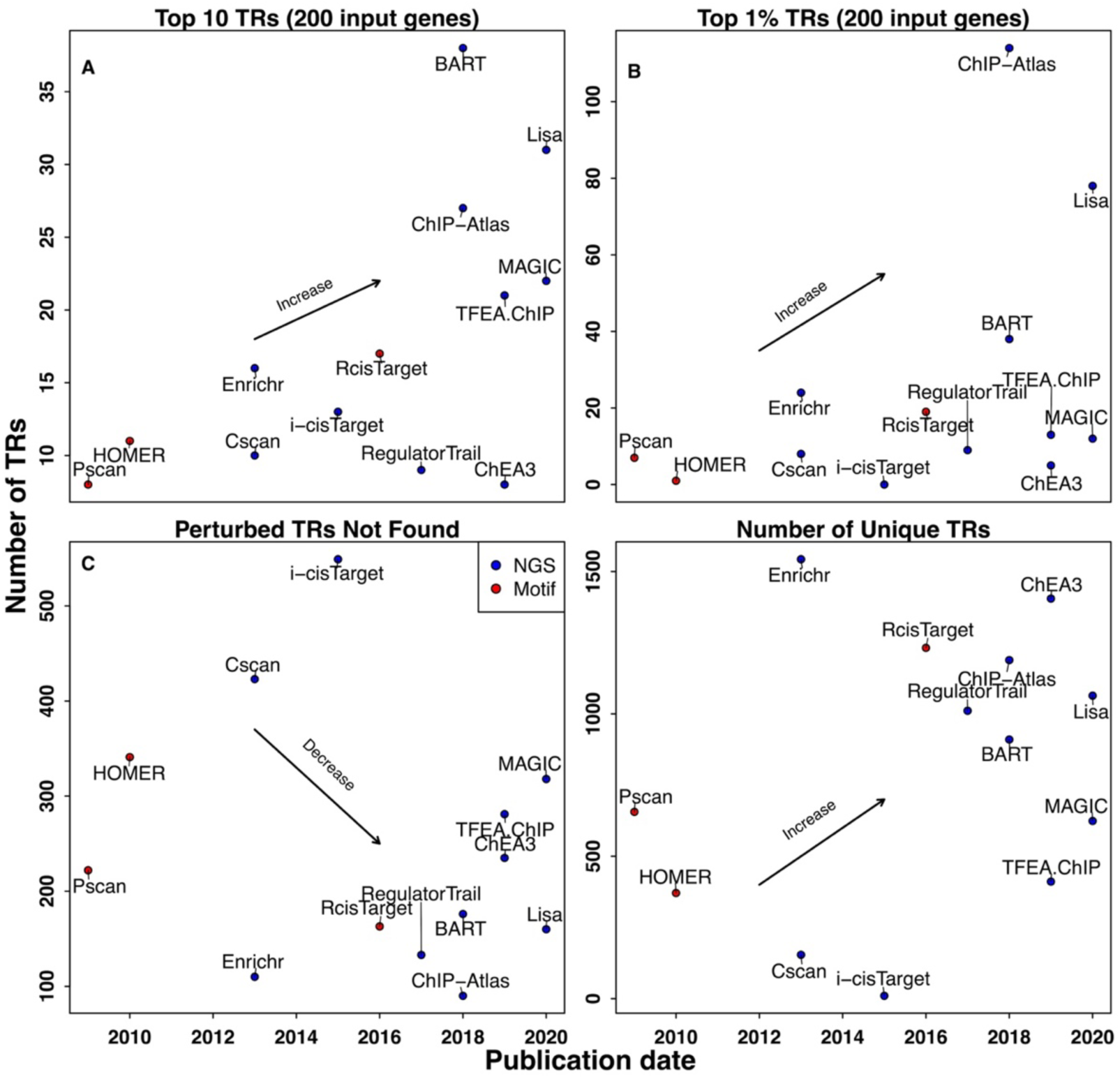
An increasing trend in prediction performance and TR coverage, aligning with more recent publication dates. (**A**) No. of perturbed TRs ranked within the top 10 positions versus method’s publication date; (**B**) No. of perturbed TRs ranked within top 1% versus method’s publication date; (**C**) No. of perturbed TRs that cannot be found in each method’s TR ranking list versus its publication date; (**D**) No. of unique TRs in the TR ranking list of each method versus its publication date.

### Usability

Figure 11 provides an assessment of NGS-based methods using various criteria of usability, with a “Yes” indicating that the criterion is satisfied and a “No” otherwise. Enrichr, TFEA.ChIP, BART, ChIP-Atlas, ChEA3, and Lisa received good scores as these methods are generally user-friendly, easy to learn and use, and did not pose any issues during testing. In summary, we recommend refraining from using methods that are not open-source, actively updated, or well-maintained. Such methods often rely on outdated databases and could pose compatibility issues.

**Figure 11.**
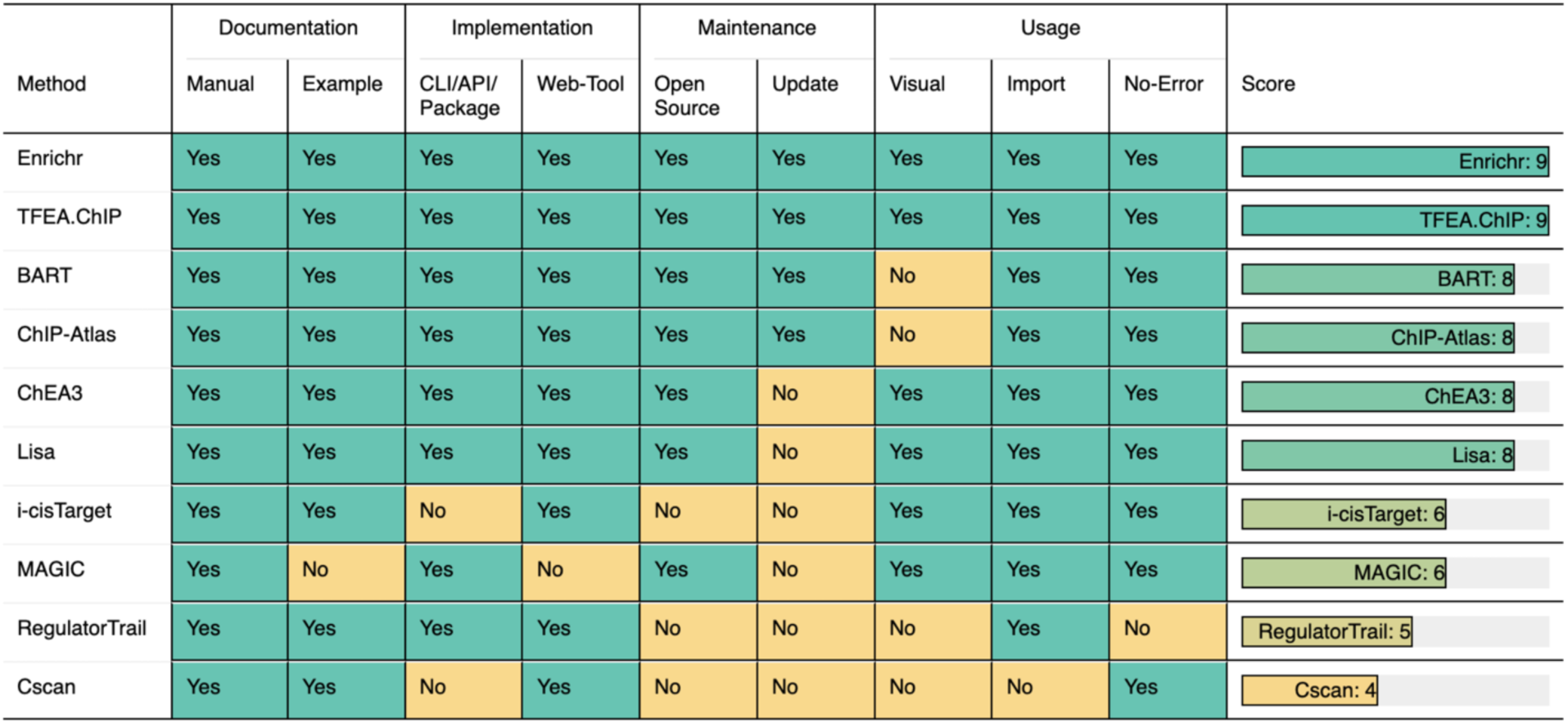
The summary of usability evaluated by using nine criteria covering most user considerations for method usage. “Yes” indicates the method satisfied the criterion and a “No” otherwise. Enrichr, TFEA.ChIP, BART, ChIP-Atlas, ChEA3, and Lisa received good scores.

## Conclusion

Our evaluation of NGS-based methods in four dimensions leads us to recommend Lisa, ChIP-Atlas, and BART, either individually or in combination, for predicting TRs of query gene sets. These methods have demonstrated relatively better performance in providing accurate predictions with perturbed TR gene sets. The evaluated methods have three major limitations in general: (1) Peaks have to be associated with genes based on linear epigenomic distance. (2) There is no ideal approach to quantify the functionality or activity of binding sites or CREs. (3) The internal database of reviewed methods can have a highly skewed distribution of the number of datasets for each TR. In addition, it is important to note that even the best-performing method may only rank less than 10% of perturbed TRs within the top 10 positions. Given the complexity of cellular processes and the limitations above, researchers should use caution when interpreting the results and should consider further validation with experimental approaches. Nevertheless, recently developed NGS-based methods provide useful tools for generating hypotheses about transcriptional regulation. Further methodological research is needed to fully utilize the wealth of NGS data and address the limitations of current methods.

## Discussion

This study evaluated the performance of various computational methods developed over the past decade for predicting TRs given a query gene set. Though NGS data can provide evidence of actual binding sites, multiple limitations make it challenging for these methods to accurately identify the functional TRs. Therefore, new experimental and computational methods are urgently needed to address these challenges. Previous studies have implicated several possible directions to improve these existing methods. First, further processing of the NGS data by using “footprinting” methods to identify binding sites in chromatin accessibility data was reported before [68]. However, this approach is limited to TRs with long residence times on chromatin, despite some TRs being known for lack of detectable footprints [69, 70]. Second, including three-dimensional long-range interaction information from Hi-C data may also serve as a possible improvement direction [71–73], but the sparse nature and low resolution of Hi-C data might also cause misleading conclusions and therefore should be used with caution and further validation.

With these limitations in mind, we also highlight that the breakthrough of single-cell sequencing techniques may lead to the development of new methods. While bulk sequencing data has dominated epigenomics research in the past decade, the limitations of bulk sequencing techniques have prompted the development of single-cell sequencing techniques such as single-cell RNA sequencing (scRNA-seq), single-cell ATAC sequencing (scATAC-seq), and single-cell ChIP-sequencing (scChIP-seq) [74]. In contrast to bulk sequencing data, these techniques sequence each cell separately and provide a more granular view of the differences between individual cells [75]. This means that limitations such as non-ideal measurement of binding affinities caused by averaging over the cells can now be addressed. In the next decade, there will likely be a surge in single-cell data, and most computational methods that aim to predict functional TRs should consider integrating single-cell data to improve their accuracy and resolution. In addition, with the rise of multi-omics experimental approaches [76, 77], the availability of multi-omics data is no longer restricted to one or two types, and there is a growing need for novel multi-omics approaches that can leverage the full range of available data. In the coming decade, it is expected that more multi-omics-based methods will be developed, which will undoubtedly lead to further innovation and discovery in this field.

## Data availability statement

Commands and scripts used to run each method can be accessed on GitHub (https://github.com/ZeyuL01/Benchmark_NGSmethods). TR perturbation derived differentially expressed genes are sourced from KnockTF [64] (https://bio.liclab.net/KnockTFv1/). ChIP-seq data are sourced from ENCODE [62] (https://www.encodeproject.org/). Top 200, 600, 1000 genes and ranking lists generated by each method are available on SMU BOX (https://smu.box.com/s/hlofza4oe08ob4bb4i02nu3wl2zy39zj).

## Funding

This work was supported by the following funding: the Rally Foundation, Children’s Cancer Fund (Dallas), the Cancer Prevention and Research Institute of Texas (RP180319, RP200103, RP220032, RP170152 and RP180805), and the National Institutes of Health funds (R21CA259771, P30CA142543, HG011996, and R01HL144969) (to L.X.)

(R01CA258584)(to X.W.)

**Zeyu Lu** is a PhD candidate in the Department of Statistics and Data Science at Southern Methodist University, who focuses on statistics and epigenomics studies.

**Xue Xiao** is a Data Scientist in the Peter O’Donnell Jr. School of Public Health at University of Texas Southwestern Medical Center, who focuses on epigenomics studies.

**Qiang Zheng** is an Assistant Professor of Research in the Department of Mathematics at University of Texas at Arlington, who focuses on statistics and epigenomics studies.

**Xinlei Wang** is Jenkins-Garrett Professor of Statistics and Data Science in the Department of Mathematics and Director of Division of Data Science for Research in College of Science, University of Texas at Arlington, who focuses on development of statistical methods and computational tools for biomedical data.

**Lin Xu** is an Assistant Professor in the Peter O’Donnell Jr. School of Public Health at University of Texas Southwestern Medical Center, who focuses on epigenomics studies.

