## Supplementary Material for "Assessing NGS-based computational methods for predicting transcriptional regulators with query gene sets"

This document includes four additional figures S1-S5 and table S1 mentioned in the main body of the paper.

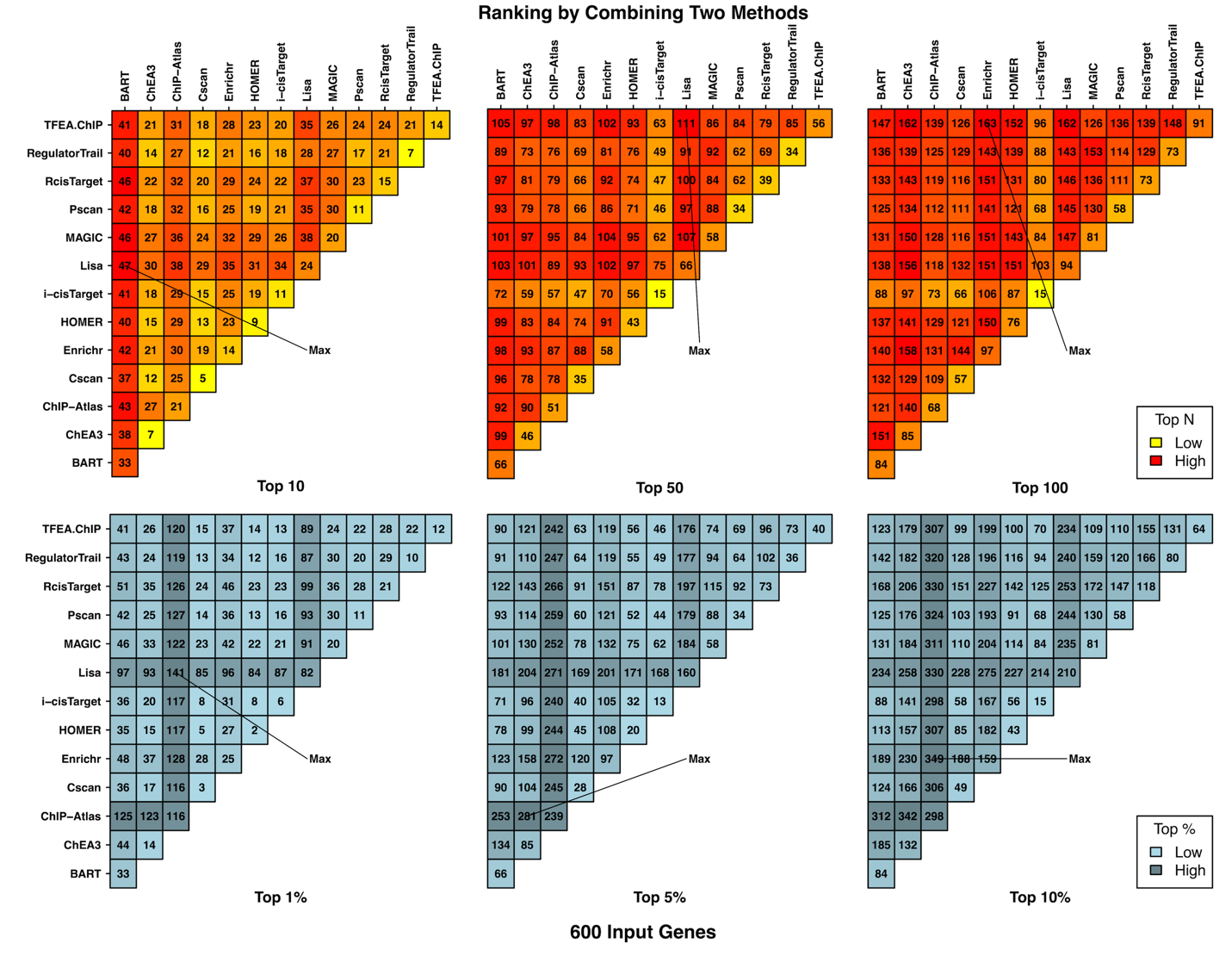

**Figure S1**. The number of perturbed TRs ranked to top positions by either method in the combination of two methods (600 input genes used). The combination of any two from BART, ChIP-Atlas, Lisa, Enrichr, and TFEA.ChIP can achieve the best or close to the best performance.

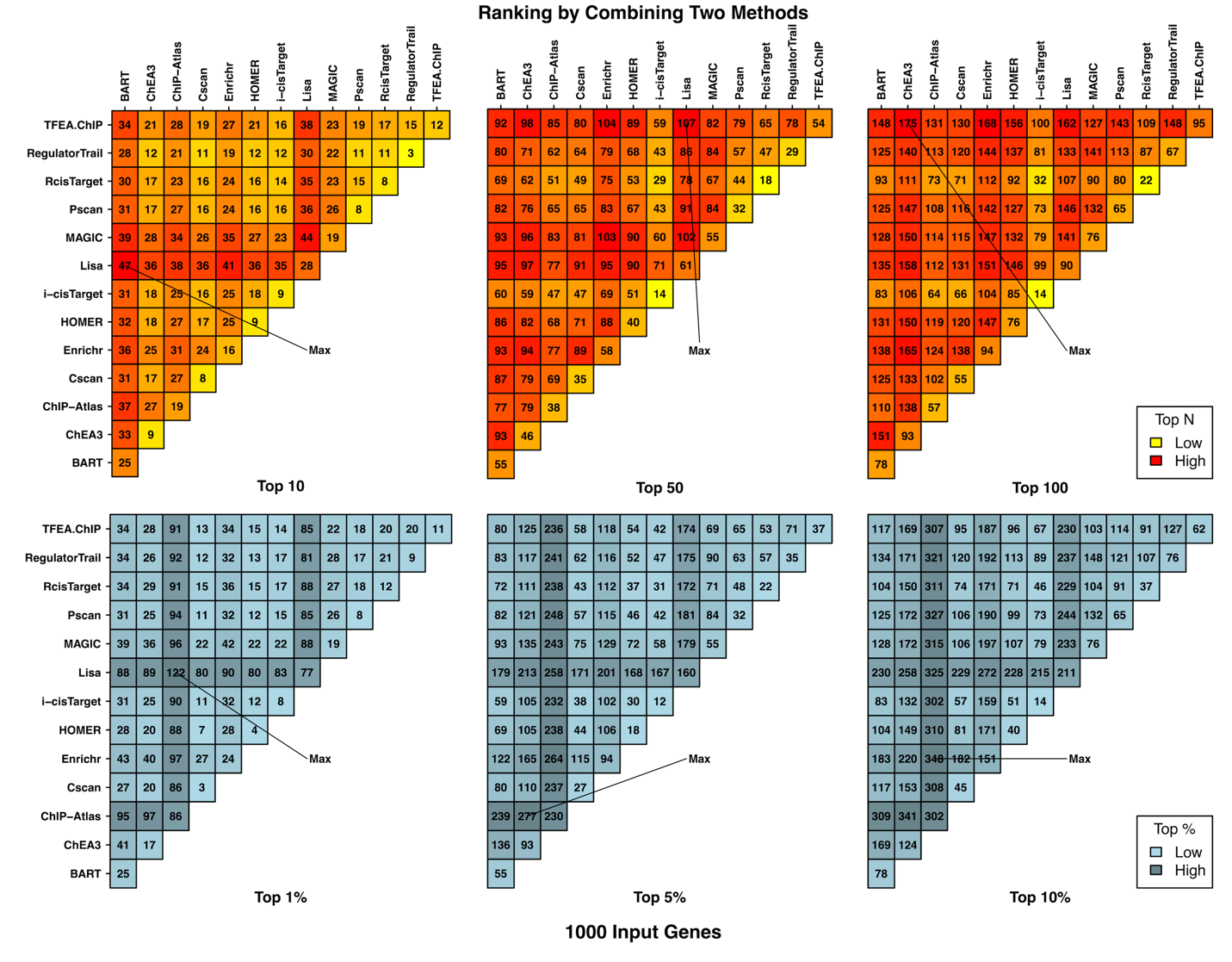

**Figure S2.** The number of perturbed TRs ranked to top positions by either method in the combination of two methods (1000 input genes used). The combination of any two from BART, ChIP-Atlas, Lisa, Enrichr, and TFEA.ChIP can achieve the best or close to the best performance.

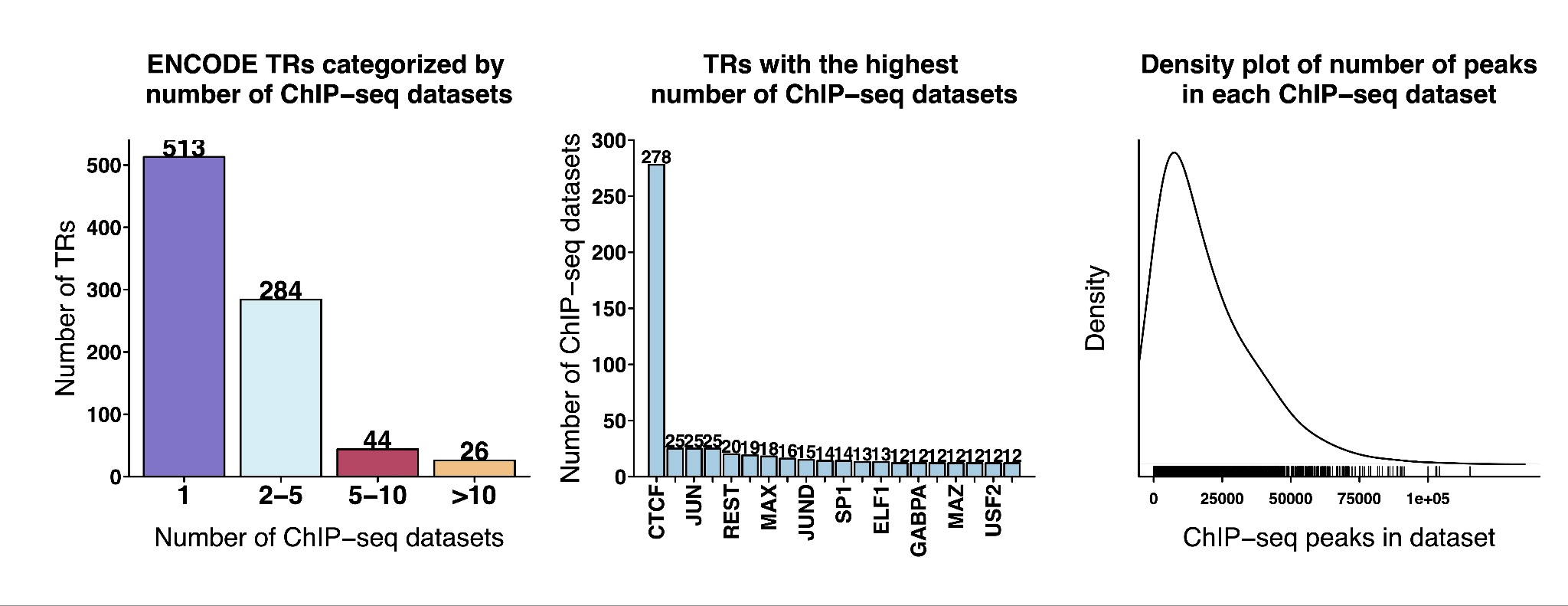

**Figure S3.** TRs can have a different number of TR ChIP-seq datasets, as shown in the ENCODE database, which is used by many methods as the main reference data resource.

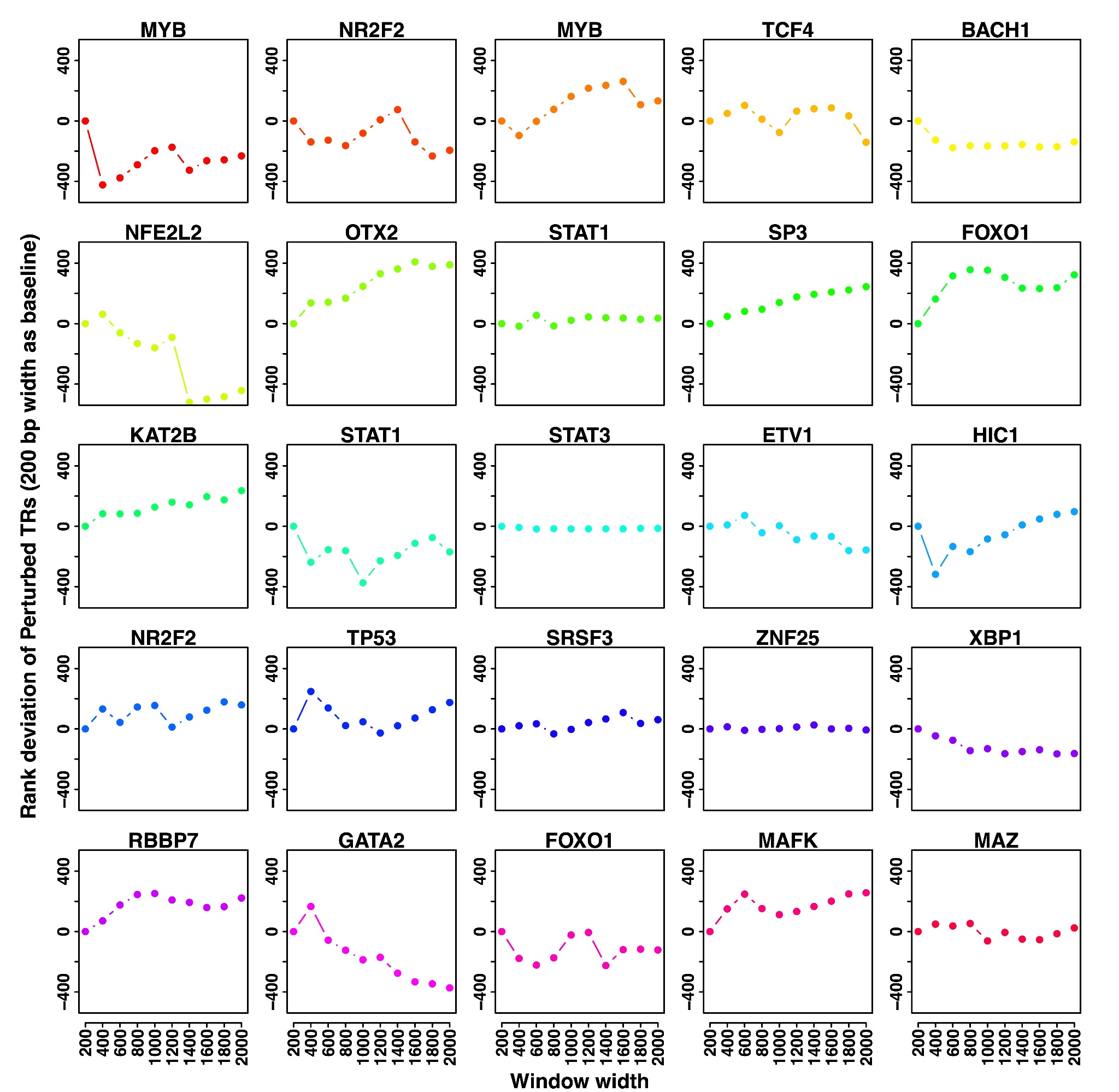

**Figure S4.** The rank deviations of 25 randomly selected perturbed TRs (using the rank for the window width of 200 bp as the baseline) by implementing a similar workflow as Cscan, a window-based approach that uses windows of pre-fixed width to assign target genes. Evidently, the ranks of some perturbed TRs are heavily affected by the cutoff choices.

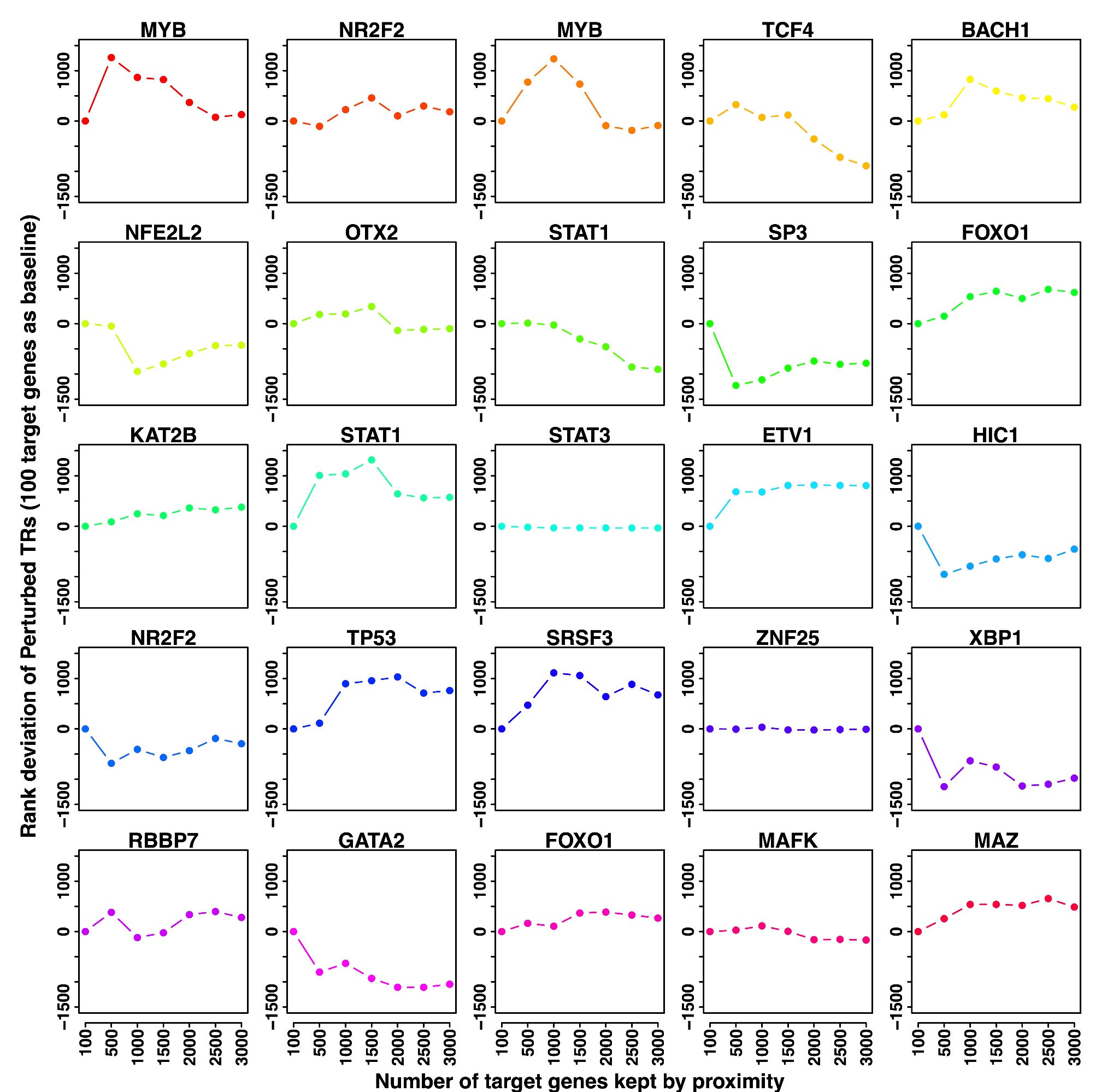

**Figure S5.** *The rank deviations of 25 randomly selected perturbed TRs (using the rank in the case of 100 target genes as the baseline) by implementing a similar workflow as Enrichr, a window-free approach that uses the number of genes kept based on the genomic distance to assign target genes. Again, the ranks of some perturbed TRs are heavily affected by the cutoff choices. There is no clear pattern of which cutoff value works better.*

| Method | Input gene set size | Stouffer's Z score | Combined P-value |
| --- | --- | --- | --- |
| BART | 200 | 13.814 | <0.001 |
|  | 600 | 16.343 | <0.001 |
|  | 1000 | 15.824 | <0.001 |
| Lisa | 200 | 87.338 | <0.001 |
|  | 600 | 82.963 | <0.001 |
|  | 1000 | 82.484 | <0.001 |
| ChIP-Atlas | 200 | 91.887 | <0.001 |
|  | 600 | 91.85 | <0.001 |
|  | 1000 | 90.576 | <0.001 |
| Cscan | 200 | 13.519 | <0.001 |
|  | 600 | 10.902 | <0.001 |
|  | 1000 | 8.876 | <0.001 |
| MAGIC | 200 | 49.670 | <0.001 |
|  | 600 | 57.646 | <0.001 |
|  | 1000 | 62.172 | <0.001 |
| TFEA.ChIP | 200 | 57.131 | <0.001 |
|  | 600 | 57.325 | <0.001 |
|  | 1000 | 58.385 | <0.001 |
| Enrichr | 200 | 60.51 | <0.001 |
|  | 600 | 52.588 | <0.001 |
|  | 1000 | 51.264 | <0.001 |
| RegulatorTrail | 200 | 102.947 | <0.001 |
|  | 600 | 110.272 | <0.001 |
|  | 1000 | 112.471 | <0.001 |
| ChEA3 | 200 | -21.898 | 1 |
|  | 600 | -18.877 | 1 |
|  | 1000 | -19.821 | 1 |

**Table S1.** We selected 20 TRs with the largest numbers of ChIP-seq datasets and 20 TRs with single ChIP-seq datasets randomly and applied the Wilcoxon rank-sum test to examine whether TRs in the first group tend to be ranked higher than those in the second group. P-values from input gene sets of the same size are combined by Stouffer’s Z scores to derive an overall p-value for each method.
